# Molybdate delays sulphide formation in the sediment and transfer to the bulk liquid in a model shrimp pond

**DOI:** 10.1101/2023.11.16.567380

**Authors:** Funda Torun, Barbara Hostins, Peter De Schryver, Nico Boon, Jo De Vrieze

## Abstract

Shrimp are commonly cultured in earthen aquaculture ponds where organic-rich uneaten feed and faeces accumulate on and in the sediment to form anaerobic zones. Since the pond water is rich in sulphate, these anaerobic conditions eventually lead to the production of sulphide. Sulphides are toxic and even lethal to the shrimp that live on the pond sediment, but physicochemical and microbial reactions that occur during the accumulation of organic waste and the subsequent formation of sulphide in shrimp pond sediments remain unclear. Molybdate treatment is a promising strategy to inhibit sulphate reduction, thus, preventing sulphide accumulation. We used an experimental shrimp pond model to simulate the organic waste accumulation and sulphide formation during the final 61 days of a full shrimp growth cycle. Sodium molybdate (5 and 25 mg/L Na_2_MoO_4_.2H_2_O) was applied as a preventive strategy to control sulphide production before oxygen depletion. Molybdate addition partially mitigated H_2_S production in the sediment, and delayed its transfer to the bulk liquid by pushing the higher sulphide concentration zone towards deeper sediment layers. Molybdate treatment at 25 mg/L significantly impacted the overall microbial community composition and treated samples (5 and 25 mg/L molybdate) had about 50% higher relative abundance of sulphate reducing bacteria than the control (no molybdate) treatment. In conclusion, molybdate has the potential to work as mitigation strategy against sulphide accumulation in the sediment during shrimp growth by directly steering the microbial community in a shrimp pond system.

## 1. Introduction

The properties of pond bottom soil (sediment) and physicochemical and microbial interactions on and in the sediment are crucial for the well-being and growth of the shrimp in aquaculture ponds (Avnimelech and Ritvo, 2003; Burford et al., 1998). Sediments contain indigenous nutrients and organic matter, derived directly from the environment, but also from uneaten and digested feed of the numerous shrimp that dwell on the pond bottom, especially during semi-intensive and intensive stocking (50-300 shrimp/m^3^) (Avnimelech and Ritvo, 2003). This pond bottom layer, *i.e.*, the interphase between the water and sediment, is an area that is densely populated by microorganisms consuming the available organic matter. Due to the organic-rich conditions on the pond bottom in combination with the typical temperatures of 25-30 °C in shrimp ponds, the oxygen consumption by these microorganisms can cause a rapid drop in dissolved oxygen in the sediment (Baxa et al., 2021; Dien et al., 2019). When oxygen consumption exceeds the rate of oxygen transfer from the pond water phase to the sediment, eventually sediment oxygen is depleted, and anaerobic conditions arise. Due to high sulphate concentrations in the pond water, low redox conditions in the pond lead to production of hydrogen sulphide (H_2_S) from metabolic activity of sulphate reducing bacteria (SRB) (Avnimelech and Ritvo, 2003; Boyd, 1998). The H_2_S formed creates a bad odour and black colour in the sediment, and is also toxic to the shrimp that dwell at the pond bottom. Sulphide toxicity to shrimp depends on both the H_2_S concentration and pH (Thulasi et al., 2020; Vismann, 1996), with lethal concentrations to kill 50% of the population (LC50) values ranging between 0.0087 and 0.033 mg/L H_2_S, depending on shrimp species and growth phase of the shrimp (Chen, 1985; US-EPA, 2011). Exposure to sub-lethal concentrations of H_2_S lowers shrimp resistance to diseases and causes tissue corrosion (Suo et al., 2017). Overall, H_2_S is often the main cause for mortality or abnormal behaviour of shrimp, and may strongly impact shrimp harvest (Panakorn, 2016).

Sulphide accumulation in shrimp ponds conventionally relies on labour-intensive, time-intensive and costly approaches, such as mechanical removal of reduced sediment or change of culture water. An alternative approach is nitrate amendment, which has been shown to remove the H_2_S produced (Torun et al., 2020). However, as also demonstrated for sodium percarbonate, the effect of nitrate towards sulphide removal was only transient, because when nitrate was depleted, the H_2_S production recovered (Schwermer et al., 2010; Torun et al., 2022; Torun et al., 2020), requiring higher amounts of nitrate addition to compete with sulphate reduction. Repeated and/or increased addition of nitrate is unwanted, because this may result in cyanobacteria and algal blooms or the release of toxic metabolites, *e.g.*, nitrite or nitrous oxide.

A more targeted, preventive approach that achieves direct inhibition of the SRB, thus, preventing sulphide production, is the application of molybdate (MoO_4_). Because of its stereochemical similarity to sulphate, molybdate inhibits the adenosine triphosphate sulfurylase, which is the first enzyme in the sulphate reduction pathway (Peck, 1959; Stoeva and Coates, 2019). Successful inhibition of sulphate reduction through the addition of molybdate has been observed in studies on eutrophic lake sediments (Smith and Klug, 1981), anaerobic digestion (Isa and Anderson, 2005; Ranade et al., 1999), and oil production systems (Jesus et al., 2015; Kögler et al., 2021). Hence, its application in aquaculture systems also warrants possibilities towards preventing H_2_S formation in pond sediments. This was demonstrated in a short-term experiment with a shrimp pond model in which molybdate outperformed nitrate and sodium percarbonate in controlling H_2_S formation, because of its specificity and preventive mode of action (Torun et al., 2022). The applicability of molybdate as a remediation strategy towards sulphide formation in aquaculture, however, strongly depends on its lasting effect during a 90-days shrimp growth cycle.

The objective of this study was to determine the duration and magnitude of the effect of molybdate towards H_2_S mitigation in response to the gradual accumulation of organic waste during a full shrimp growth cycle. A shift in the microbial community towards different processes than sulphate reduction, in response to molybdate, could be beneficial to control sulphide accumulation at the shrimp pond bottom. Because the accumulation of organic waste was limited during the first 30 days of the shrimp growth cycle, *i.e.*, no O_2_ depletion or H_2_S accumulation (Torun et al., 2023), only the final 61 days were considered in a lab-scale shrimp pond bottom model.

## 2. Material and methods

### 2.1. Sampling and storage

#### 2.1.1. Sediment sampling

Sandy clay and organic-rich sediments were obtained from the Ijzermonding Nature Reserve (Nieuwpoort, Belgium) from a creek (51°8′45′′ N/2°44′38′′E) that was regularly water-logged with tidal movement. Sampling were taken by scooping the top 5-10 cm of the sediment into a closed plastic container in which they were transported to the laboratory. The pH, conductivity, total solids (TS) and volatile solids (VS) of the fresh sediment were analysed directly upon arrival in the laboratory. A sample for sulphate and molybdate analysis was stored at 4°C until analysis, and a sample for DNA extraction was stored at −20°C.

#### 2.1.2. Feed and faeces collection and storage

Fresh shrimp faeces were collected from the flush outlet of shrimp tanks in which whiteleg shrimp (*Litopenaeus vannamei*) at post-larvae stage were fed with CreveTec Grower 2 (CreveTec, Ternat, Belgium) at the Aquaculture and *Artemia* Reference Center (ARC), Faculty of Bioscience and Engineering, Ghent University, Belgium. The faeces were stored at 4°C until use to avoid organic matter degradation during storage. The pH and conductivity of the faeces were measured directly after collection. A sample for DNA extraction was stored at −20°C.

### 2.2. Experimental set-up and operation

A system mimicking organic matter accumulation in the shrimp ponds was designed and constructed, as described earlier (Torun et al., 2022), using 250 mL size glass beakers (outer diameter 70 mm) containing a 3.5 cm sediment layer and 5 cm overlaying artificial seawater (Instant Ocean, Aquarium Systems, Mentor, OH, USA). The salinity of the artificial seawater was adjusted to 20 g/L, representing a common salinity in shrimp ponds (15-25 g/L), and containing approximately 1.5 g/L sulphate. To avoid excessive water evaporation, the beakers were put in a transparent plastic box with a non-airtight lid in a temperature-controlled room at 28 ± 1°C without active aeration. No artificial or natural light was foreseen to avoid the growth of microalgae and keep a focused approach towards sulphide formation and oxygen depletion.

The experiment was started with an initial cumulative waste of 30 days of shrimp culture (DOC 30) in the form of feed (CreveTec Grower 2 shrimp feed) and faeces, which was considered day zero of the experiment. The cumulative waste for shrimp culture was calculated based on the 0.003848 m^2^ bottom surface area of the beakers using commercial daily shrimp feeding tables (Table S1). Feed and faeces were added based on semi-intensive stocking of 50 shrimp/m^2^. About 25% of input feed was assumed to be accumulating in the pond bottom, with 15% considered digested feed (faeces), and 10% as uneaten feed. After adding the initial waste of DOC 30, the respective amount of shrimp feed and faeces, based on daily uneaten feed and faeces, were supplemented every 2-3 days (Table S2). The amounts of supplemented feed and faeces were increased every 15 days to adapt to the growth of the shrimp.

Two different concentrations of 5 (M5) and 25 (M25) mg/L of sodium molybdate (Na_2_Mo_4_.2H_2_O, Sigma Aldrich, St. Louis, Mo., US), were compared with a control treatment (no molybdate addition) for the last 61 days of a shrimp growth cycle. Molybdate was supplemented in a single dose on day 0 of the experiment. Each treatment was carried out in 6 biological replicates. Measurements of dissolved oxygen (DO), H_2_S and pH in the bulk liquid were performed every 2-3 days. These measurements were taken from the water column, about 1 cm above the sediment-water interface, since H_2_S in the bulk liquid is the major concern for the shrimp that dwell on the pond bottom. Apart from bulk liquid measurements, microscale gradient depth profiles of DO, H_2_S and pH at the water-sediment interface and throughout the sediment were measured, using a microelectrode, on day 16, 30, 44 and 61 from three replicates of each treatment. After each depth profiling measurement (day 16, 30, 44), one replicate from each treatment was sacrificed for the measurement of molybdate and sulphate concentrations in the bulk liquid. From these sacrificial beakers, sediment samples were taken, and stored at −20°C for microbial community analysis. At the end of the experiment (day 61), all remaining replicates (3 replicates) were sampled for sediment and liquid samples. Sediment samples were taken from the upper 1 cm of the sediment layer after carefully decanting the liquid part. The liquid samples were filtered over a 0.20 μm Chromafil^®^ Xtra filter (Macherey-Nagel, PA, USA), and stored at 4 °C, prior to analysis of sulphate and molybdate concentrations. During each liquid sampling, the degree of water evaporation was determined by recording the water depth. The molybdate and sulphate measurements were corrected with the evaporation factor.

### 2.3. Microelectrode measurements

Microscale depth profiles of O_2_, pH and H_2_S were recorded using commercial microelectrodes (Unisense A.S. Denmark, tip sizes pH: 200 μm, H_2_S: 100 μm, O_2_: 100 μm), operated with a motorized micromanipulator (Unisense A. S., Denmark). Microscale measurements were always performed before other samples were taken and before adding fresh waste to avoid disturbance of the water column and sediment. The oxygen profiles were measured at 200 μm resolution. The pH and H_2_S were simultaneously recorded with the same resolution at 200 μm in the water-sediment interphase, and at lower resolution deeper in the sediment. The sensors were calibrated following standard calibration procedures, as described earlier (Malkin et al., 2014). The H_2_S was calibrated with a 3-5 point standard curve using an acidified Na_2_S standard solution (pH 3.5-4.0). The O_2_ sensor was calibrated with a 2 point standard curve, using 100% in air bubbled seawater for the DO at saturation at 28°C and argon bubbled seawater for DO zero. The pH sensor was calibrated with 2 point calibrations using commercial (Carl Roth GmbH & Co.KG, Karlsruhe, Germany) pH buffer solutions (4, and 7). Total sulphide concentrations were calculated as described earlier (Jeroschewski et al., 1996). For bulk liquid measurements, the same electrodes were used manually to take the measurements from approximately 1 cm above the sediment surface after ensuring that there was negligible variety in the duplicate measurements of the water column parameters.

### 2.4. Analytical techniques

The TS and VS of the sediment were determined according to Standard Methods (Greenberg et al., 1992). The pH of the overlaying water and sediment samples were measured with a pH meter (Metrohm, Herisau, Switzerland), which was calibrated using pH buffer solutions at pH 4 and 7. The sulphate concentrations were measured through ion chromatography (930 Compact IC Flex, Metrohm, Herisau, Switzerland), equipped with a Metrosep A supp 5– 150/4.0 anion column with conductivity detector, after diluting the samples 1:50 using ultra-pure water (Milli-Q, Millipore Corporation, Burlington, MA, USA). The detection range was 0.05 to 200 mg ion/L. Molybdate was measured using a commercial kit (Hach, Model Mo-2, USA), based on the colorimetric determination of molybdenum using mercaptoacetic acid (Will and Yoe, 1953). Standard solutions of 0, 5, 10, 25, and 50 mg/L Na_2_MoO_4_.2H_2_O were prepared to determine the standard curve at 425 nm using a UV-Vis Spectrophotometer (WPA Lightwave II, Thermofisher, USA).

### 2.5. Microbial community analysis

#### 2.5.1. Amplicon sequencing

To analyse the changes in the bacterial community and SRB relative abundance, samples were taken from the upper 1 cm of the sediment from each sacrificial beakers in 3 replicates and frozen at −20 °C. The DNA was extracted directly from the frozen samples using a commercial kit (DNeasy Power Soil Pro Kit, QIAGEN, Hilden, Germany), following the instructions of the manufacturer. The quality of the DNA extracts was evaluated through agarose gel electrophoresis and PCR analysis with the universal bacterial primers 341F (5’-CCTACGGGNGGCWGCAG) and 785Rmod (5’-GACTACHVGGGTATCTAAKCC) that target the V3-V4 region of the 16S rRNA gene (Klindworth et al., 2013), following a PCR protocol as described earlier (Boon et al., 2002). The samples were sent to LGC Genomics GmbH (Berlin, Germany) for Illumina amplicon sequencing of the V3-V4 region of the 16S rRNA gene of the bacterial community on the MiSeq platform with V3 chemistry. The amplicon sequencing protocol and data processing are described in detail in the SI (S3).

#### 2.5.2. Flow cytometry analysis

Absolute microbial cell counts in the sediment samples were determined using flow cytometry (FCM). Prior to FCM analysis, sediment samples were defrosted, acclimated to room temperature and diluted tenfold in sterile, 0.22 µm-filtered Instant Ocean® solution. To separate the cells from sediment particles, samples were initially sonicated (Q700 Sonicator, Qsonica, Newtown, CT, USA) for 3 minutes, followed by 3 minutes centrifugation at 500 *g*. The resulting supernatant of the samples was stained with 1 vol% SYBR® Green I (SG, 100x concentrate in 0.22 µm-filtered DMSO, Invitrogen), and incubated in the dark at 37°C for 20 min. Immediately after incubation, samples were analysed using a BD Accuri C6 Plus cytometer (BD Biosciences, Erembodegem, Belgium), equipped with four fluorescence detectors (533/30 nm, 585/40 nm, > 670 528 nm and 675/25 nm), two scatter detectors and a 20-mW 488-nm laser. Samples were analysed in fixed volume mode (30 µL). The flow cytometer was operated with Milli-Q water (MerckMillipore, Belgium) as sheath fluid, and instrument performance was verified daily using CS&T RUO Beads (BD Biosciences, Erembodegem, Belgium).

### 2.6. Statistical analysis

A table containing the relative abundances of the different OTUs (operational taxonomic units), and their taxonomic assignment was created following amplicon data processing (Supplementary Information File 2). All statistical analysis were carried out in R Studio version 4.03 (http://www.r-project.org) (R Development Core Team, 2013). Absolute singletons were removed, and the different samples were rescaled *via* the “common-scale” approach (McMurdie and Holmes, 2014) through which the proportions of all OTUs were taken, multiplied with the minimum sample size, and rounded to the nearest integer. Sampling depth of each sample was evaluated through rarefaction curves (Figure S1) (Hurlbert, 1971; Sanders, 1968). The packages vegan (Oksanen et al., 2016) and phyloseq (McMurdie and Holmes, 2013) were used for microbial community analysis. A heatmap was created at the phylum and family level (1% cut-off) with the *pheatmap* function (pheatmap package), and biological replicates were collated according to the method described earlier (Connelly et al., 2017). The non-metric multidimensional scaling (NMDS) plots were constructed using the Bray-Curtis (Bray and Curtis, 1957) distance measures. Significant differences between treatments and timepoints were identified using pairwise permutational ANOVA (PERMANOVA) analysis (9999 permutations) with Bonferroni correction, using the *adonis*function (vegan).

## 3. Results

### 3.1. Impact of molybdate on oxygen depletion and sulphide production

#### 3.1.1. Bulk liquid concentrations

The DO measurements in the bulk liquid showed that oxygen was completely depleted for the first time on day 7 for all treatments (Figure 1a). In the following days, there was a fluctuation in oxygen concentrations, *i.e.*, between 0 and 150 µM, for all treatments, with a regular oxygen re-introduction in the bulk between day 7 to 35, keeping in mind that no active aeration was applied. After day 35 of the experiment, a full oxygen depletion was observed for all treatments with only limited oxygen re-introduction, resulting in oxygen concentrations only up to 50 µM. For the entire experimental period, the DO concentration in the bulk did not show clear difference between the different treatments, and also pH remained similar in the different treatments (Figure S2).

**Figure 1.**
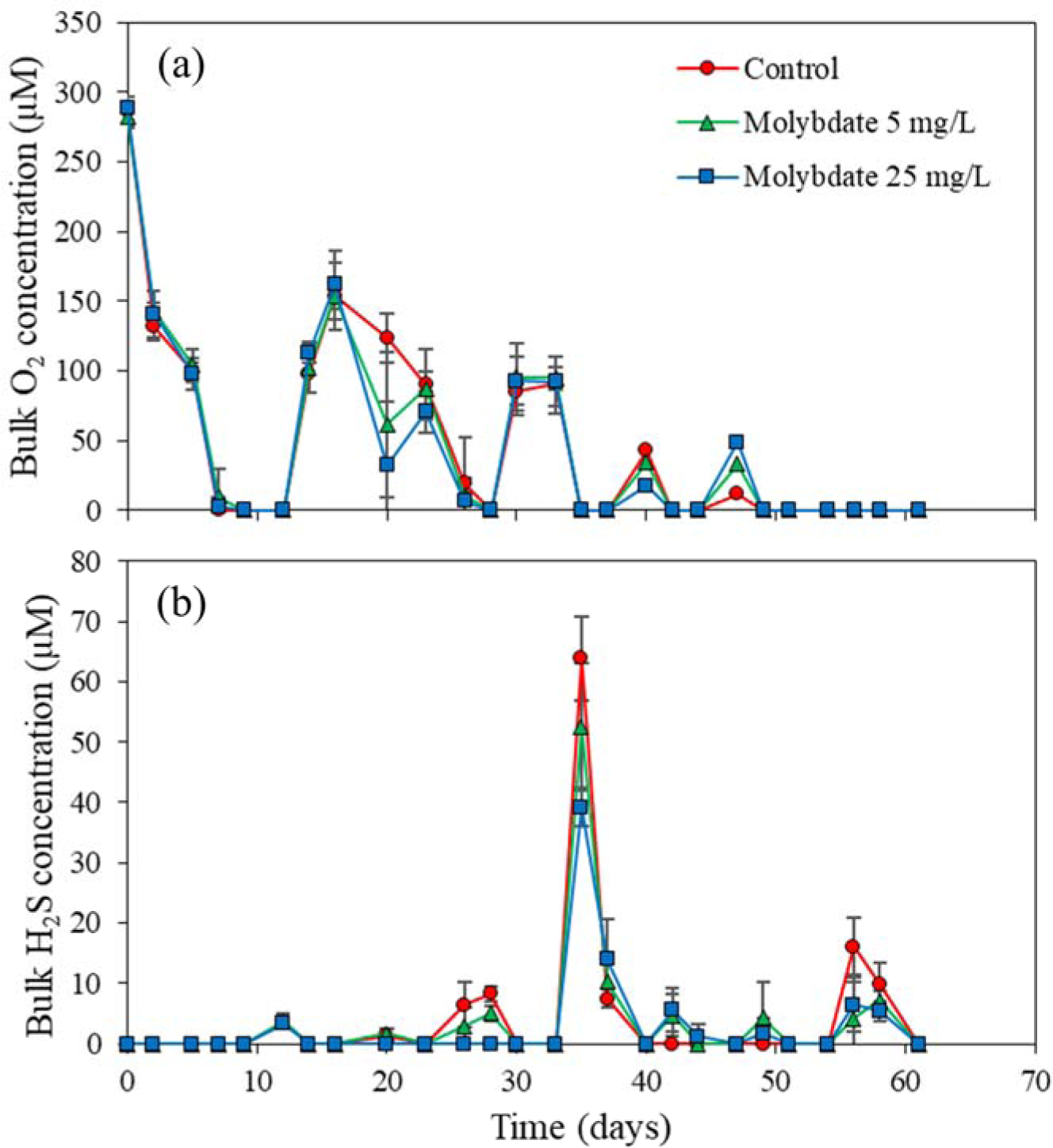
The bulk liquid concentrations of (a) O_2_ and (b) H_2_S in the control treatment, molybdate treatment at 5 mg/L (M5) and molybdate treatment at 25 mg/L (M25). Values represent averages of biological triplicates, and error bars represent the standard deviation.

No clear H_2_S production was observed, *i.e.*, H_2_S concentrations did not exceed 10 µM, in the bulk liquid until day 35 of the experiment, coinciding with the time when oxygen depletion for all incubations was recorded (Figure 1b). On day 35, a H_2_S concentration of 64 ± 7, 53 ± 11, and 39 ± 3 µM was recorded in the bulk liquid for the control, M5, and M25 treatments, respectively. This corresponded with a total bulk sulphide concentration that was 22 ± 1 % and 46 ± 1 % lower than the control for M5 and M25, respectively (Figure S3). Hence, there was markedly lower H_2_S production in the molybdate treatments compared to the control treatment, especially in the M25 treatment. Also on day 56 of the experiment, H_2_S concentration in the bulk was clearly higher in the control treatment (16 + 5 µM) compared to the M5 (7 + 6 µM) and M25 (5 + 5 µM) treatments.

Residual molybdate measurements indicated that about 53 ± 1 % of the dosed molybdate disappeared from the bulk liquid for M25 at the end of the experiment, while this ranged between 5-15% for M5. (Table 1). Hence, in both molybdate treatments, residual molybdate remained present. Residual sulphate concentration in the bulk gradually decreased throughout the experiment, yet, no clear differences could be observed between the treatments (Table 2).

**Table 1.** Molybdate concentrations of bulk liquid samples taken from the sacrificed replicates on day 16, 30 and 44. At the end of the experiment (day 61), all three remaining replicates were analysed, hence, the values for day 61 are average values and standard deviations of biological replicates. M5 = Treatment with 5 mg/L molybdate addition. M25 = Treatment with 25 mg/L molybdate addition.

|  | Molybdate concentration (mg/L) |  |  |  |
| --- | --- | --- | --- | --- |
|  | Day 16 | Day 30 | Day 44 | Day 61 |
| Control | 0.0 | 0.0 | 0.0 | 0.0 ± 0.0 |
| M5 | 5.9 | 4.2 | 4.5 | 4.8 ± 4.3 |
| M25 | 21.9 | 15.7 | 12.0 | 11.6 ± 0.7 |

**Table 2.** Sulphate concentrations of bulk liquid samples taken from the sacrificed replicates on day 16, 30 and 44. At the end of the experiment (day 61), all three remaining replicates were analysed, hence, the values for day 61 are average values and standard deviations of biological replicates. M5 = Treatment with 5 mg/L molybdate addition. M25 = Treatment with 25 mg/L molybdate addition.

|  | Sulphate concentration (mg/L) |  |  |  |
| --- | --- | --- | --- | --- |
|  | Day 16 | Day 30 | Day 44 | Day 61 |
| Control | 1396 | 1207 | 1283 | 1170 ± 206 |
| M5 | 1334 | 1201 | 1261 | 1021 ± 75 |
| M25 | 1389 | 1206 | 1351 | 1100 ± 78 |

#### 3.1.2. Sediment profiles

Microscale depth profiles of the DO in the sediment on day 16 revealed a 12 ± 4 % higher concentration of oxygen in the M25 compared to the control treatment at the water-sediment interface (Figure 2). Oxygen diffusion into the first 2 mm of the upper sediment layer was observed for all treatments, with negligible differences between the treatments. On day 30, the DO depth profile showed no apparent difference between the different treatments On day 44 and 61, no more oxygen was detected in the bulk liquid or sediment.

**Figure 2.**
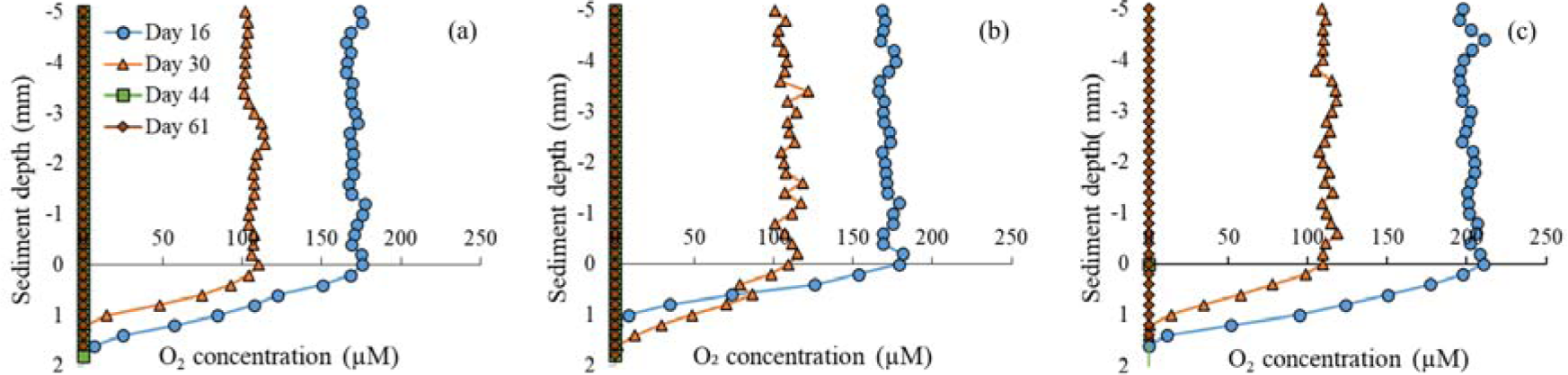
The O_2_ depth profiles for the (a) control treatment, (b) molybdate treatment at 5 mg/L (M5) and (c) molybdate treatment at 25 mg/L (M25). Values represent averages of biological triplicates, error bars are omitted to maintain the visibility of the graphs. Zero depth equals to the sediment-water interface. On day 44 and 61, all O_2_ values were below the detection limit.

Microscale depth profiles of H_2_S in the sediment were recorded on day 16, 44 and 61, while day 30 gradient measurements of H_2_S could not be obtained, due to technical problems with the microelectrode (Figure 3). On day 16, although no H_2_S could be observed in the bulk liquid, H_2_S gradient measurements showed a minor H_2_S production in the control reaching a concentration up to 5.8 ± 0.1 µM at the sediment depth of 3.6 mm, while for the M5 and M25 treatments, no H_2_S production was observed. On day 44, the control treated showed a maximum H_2_S concentration of 66.6 ± 20.5 µM in the sediment deeper layers (sediment depth of 6.0 mm). The M5 and M25 treatments showed a maximum H_2_S concentration of only 22.7 ± 4.2 µM and 29.3 ± 26.8 µM H_2_S, respectively, both at 7.2 mm sediment depth, being 69 ± 11 and 60 ± 39 % lower than the maximum value in the control treatment, respectively. On day 61, the H_2_S concentration in the sediment was more similar between the different treatments, in contrast to day 16 and 44. The M5 and M25 treatments showed a 17 ± 9 % and 26 ± 15 % lower maximum H_2_S concentration compared with the control, respectively. The H_2_S production zone appeared to be pushed to deeper sediment layers in the M5 and M25 treatments both for day 44 and 61 measurements, compared to the control. Total S microscale depth profiles showed a similar pattern as the H_2_S profiles (Figure S4), with the M5 and M25 treatments showing a markedly lower total S concentration in the sediment in comparison with the control treatment. The pH microscale depth profiles were similar between the different treatments, with limited variation in function of time (Figure S5).

**Figure 3.**
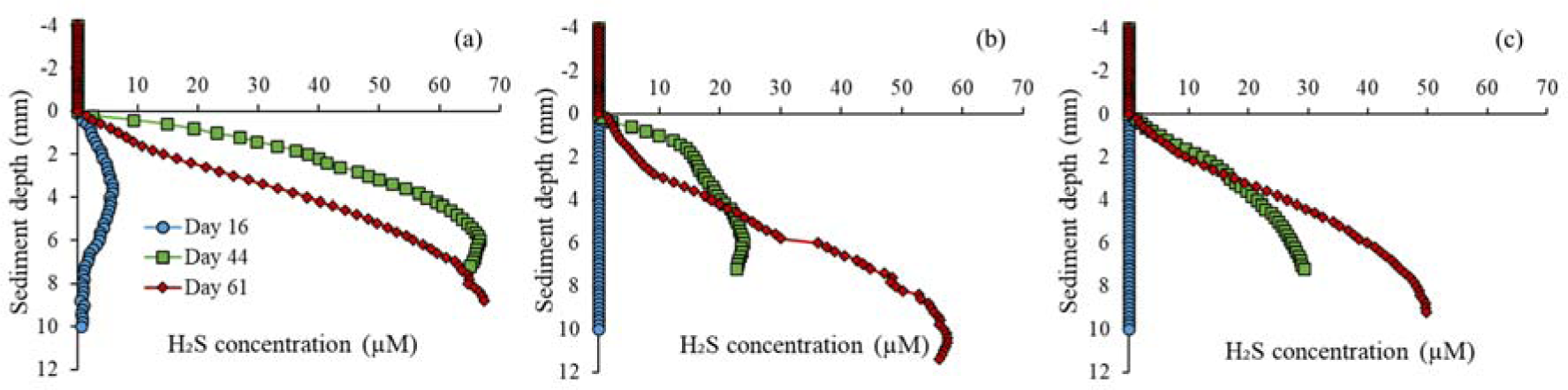
The H_2_S depth profiles for the (a) control treatment, (b) molybdate treatment at 5 mg/L (M5) and (c) molybdate treatment at 25 mg/L (M25). Values represent averages of biological triplicates, error bars are omitted to maintain the visibility of the graphs. Zero depth equals to the sediment-water interface. Because of technical problems with the microelectrode, data from day 30 are not included.

### 3.2. Microbial community analysis

Amplicon sequencing of the bacterial community resulted in an average of 23,917 ± 9,397 reads, which represented 2,922 ± 876 OTUs per sample (including singletons). Removal of absolute singletons and rescaling through the “common-scale” approach resulted in an average of 6,031 ± 282 reads and 681 ± 171 OTUs per sample.

Shrimp faeces (day 0) were dominated by Bacteroidota (35.9 ± 7.8 %) and Fusobacteriota (35.1 ± 5.7%) phyla, while the sediment used in the experiment was dominated by Actinobacteriota (22.3 ± 8.8%) and Proteobacteria (33.5 ± 1.6%) phyla (Figure 4). The sediment samples (day 0) showed a relatively higher abundance of the phylum Desulfobacterota (4.2 ± 0.3%), which contains several sulphate reducers, than the shrimp faeces (0.2 ± 0.0 %) samples. Over time, there was a clear shift in the bacterial communities, specifically for Proteobacteria, Bacteroidota and Desulfobacterota relative abundances for all treatments. On day 16, the control, M5 and M25 treatments showed markedly high relative abundances of Proteobacteria (29.3 ± 3.2 %, 27.2 ± 5.8 %, 30.7 ± 0.4 %, respectively). On day 30 and 44, the M5 and M25 treatment showed an even further increase in the relative abundance of Proteobacteria (35.6 ± 6.7 % and 34.7 ± 6.2 % for day 30 and 34.2 ± 8.1% and 38.0 ± 0.2 % for day 44, respectively), compared to the control treatment (26.6 ± 4.0 % for day 30, 22.8 ± 3.1 % for day 44). This higher relative abundance of Proteobacteria in the M5 and M25 treatments coincided with a more prominent presence of the Rhodobacteraceae family (Figure S6) in the M5 (16.1 ± 4.5 %) and M25 (14.5 ± 7.2 %) treatments, compared to the control (8.3 ± 0.6%), on day 30 and later timepoints in the experiment. The M5 (15.3 ± 3.2 %) and M25 (16.1 ± 4.1 %) treatment showed an overall higher relative abundance of Flavobacteriaceae than the control treatment (12.3 ± 2.7 %). The high relative abundance of Bacteroidota (Figure 4) likely originated from the addition of faeces, and reached values of 27.9 ± 1.2 %, 27.9 ± 0.8 % and 20.5 ± 1.8 % on day 16 for the control, M5 and M25 treatments, respectively. However, in time, the relative abundance of Bacteroidota decreased in all treatments (22.3 ± 2.4%). There was no clear difference in Bacteroidota relative abundance between the different treatments.

**Figure 4.**
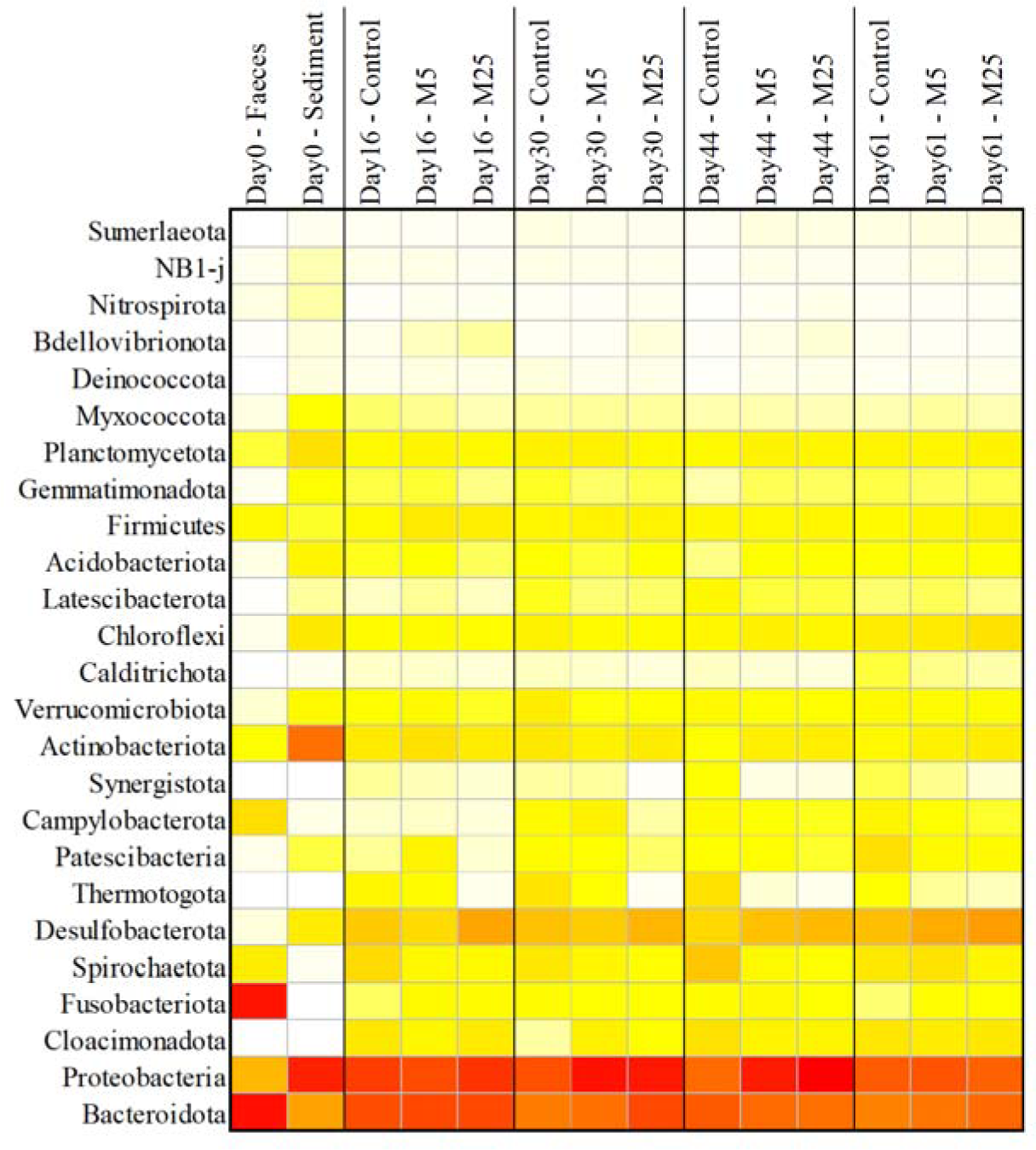
Heatmap showing the relative abundance of the bacterial community at the phylum level in the faeces, the sediment and the different treatments on day 16, 30, 44 and 61. Weighted average values of the biological replicates are presented. The colour scale ranges from 0 (white) to 40% (red) relative abundance.

There was an apparent higher abundance of the Delsulfobacterota phylum, which contains several sulphate reducers, in the M5 (15.3 ± 3.1 %) and M25 (16.2 ± 0.4%) treatments, compared to the control treatment (8.8 ± 1.6%) for all samples on day 16, 30, 44 and 61. The M25 treatment also showed a slightly higher abundance of this phyla compared to the M5 treatment. Family level analysis revealed that SRB species belonged to Desulfobulbaceae, Desulfomonadaceae, Desulfolunaceae, Delsufovibrionaceae, Desulfosarcinaceae, Delsulfobacteraceae and Desulfocapsaceae families (Figure S6). An absolute cell count analysis of Delsulfobacterota phylum, by combining flow cytometry cell counts with amplicon sequencing data, showed that all samples, including the samples treated with molybdate, showed an increasing trend in time of absolute Delsulfobacterota cell counts. Molybdate treated samples, especially M25, in general, showed even higher absolute cell counts for the Desulfobacterota phylum compared to the control (Table 3).

**Table 3.**
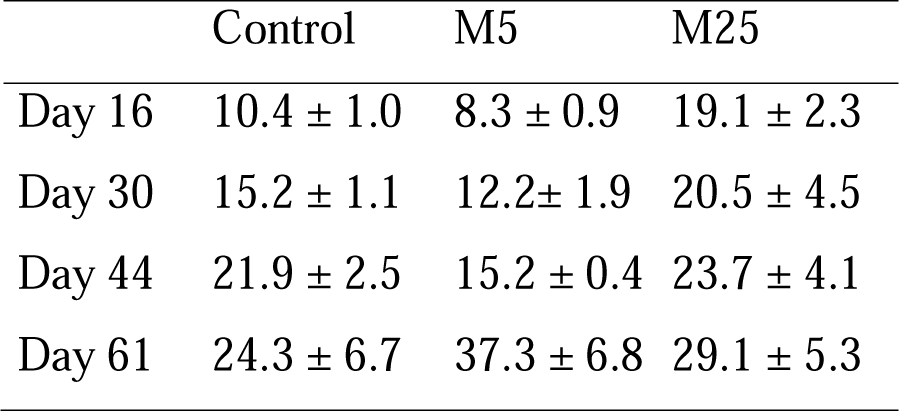
Absolute cell counts of the Desulfobacterota (10^4^ cells per mL), which contains several sulphate reducing bacteria (SRB), as determined by combing flow cytometry cell counts with amplicon sequencing data. The values are average values and standard deviations of biological replicates. M5 = Treatment with 5 mg/L molybdate addition. M25 = Treatment

|  | Control | M5 | M25 |
| --- | --- | --- | --- |
| Day 16 | $10.4 \pm 1.0$ | $8.3 \pm 0.9$ | $19.1 \pm 2.3$ |
| Day 30 | $15.2 \pm 1.1$ | $12.2 \pm 1.9$ | $20.5 \pm 4.5$ |
| Day 44 | $21.9 \pm 2.5$ | $15.2 \pm 0.4$ | $23.7 \pm 4.1$ |
| Day 61 | $24.3 \pm 6.7$ | $37.3 \pm 6.8$ | $29.1 \pm 5.3$ |

The β-diversity analysis of the bacterial community, based on the Bray-Curtis distance measure, revealed that the M25 treatment showed an overall significantly different bacterial community composition than the control treatment (*P* = 0.0003) (Figure 5). However, none of the other treatments significantly differed (*P* > 0.05), and there was a limited impact of molybdate addition on the change of the bacterial community in function of time. The PERMANOVA analysis showed that there was a significant change in overall bacterial community composition between day 16 and 30 (*P* = 0.0036), day 30 and 44 (*P* = 0.0174) and day 44 and 61 (*P* = 0.0066). On day 61, at the end of the experiment, the bacterial community composition for all treatments showed a clear divergence.

**Figure 5.**
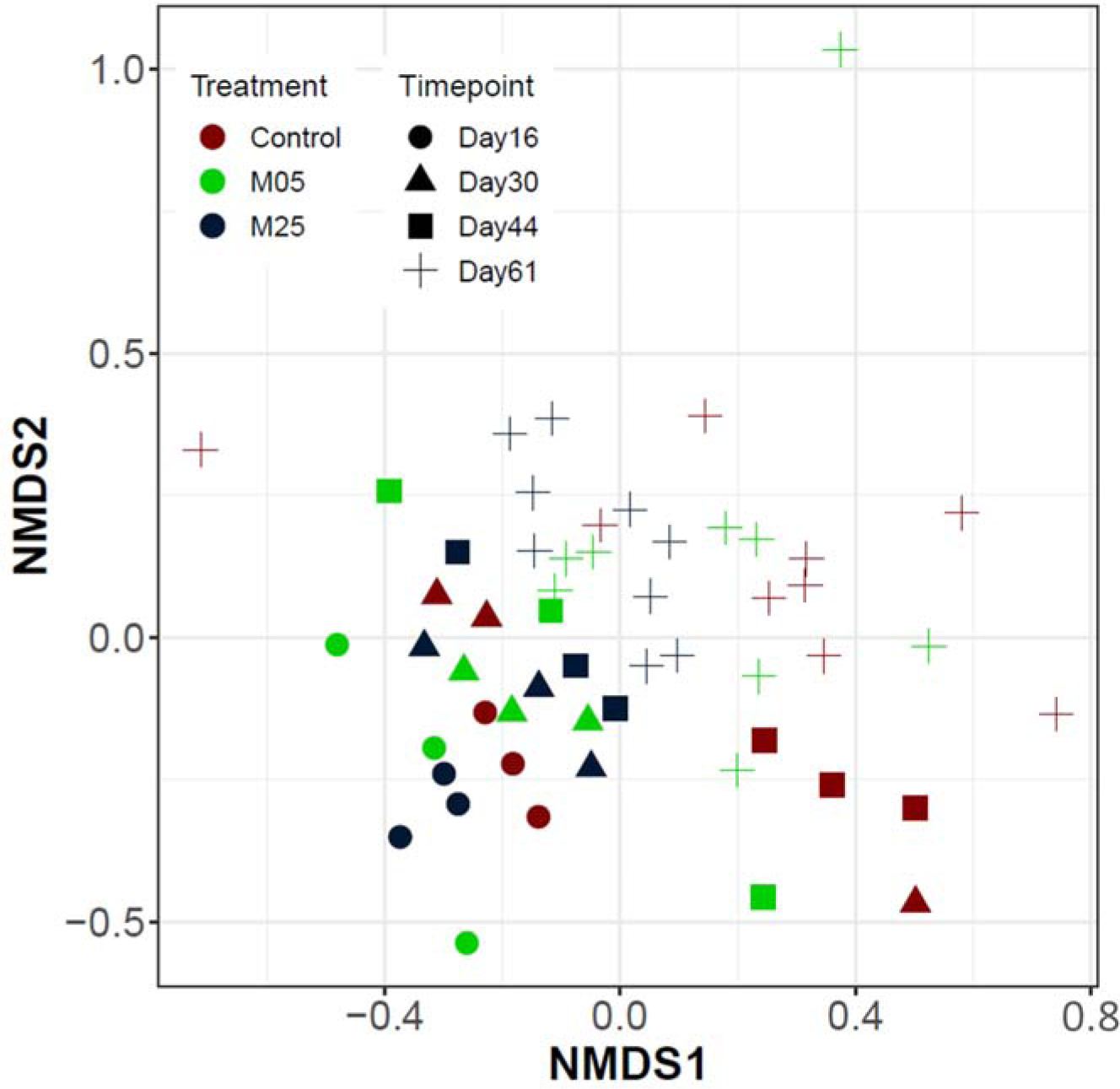
Non-metric multidimensional distance scaling (NMDS) analysis of the Bray-Curtis distance measure of the bacterial community based on amplicon sequencing data at OTU level. Different colours and symbols are used for different treatments and timepoints, respectively.

## 4. Discussion

This study showed that molybdate addition, prior to H_2_S formation, has a good potential to mitigate H_2_S production in the sediment, and delay its transfer to the bulk liquid by pushing sulphide production zone in deeper layers of the sediment. Bacterial community analysis revealed a limited impact of molybdate addition on the change of the bacterial community in function of time. Molybdate treated samples did show a higher absolute abundance of the Desulfobacterota phylum compared to the control.

### 4.1. Molybdate effectively controls sulphide production and pushes higher sulphide concentration zones towards deeper sediment layers

The typical shrimp pond water with a salinity of 1.5-2.5 % contains about 1500 mg/L sulphate (Torun et al., 2020). This high availability of sulphate and organic-rich conditions in the pond bottom make the shrimp pond environment susceptible to the production of sulphides when anaerobic conditions arise, due to the depletion of oxygen. The most effective method for avoiding anaerobic conditions is to keep dissolved oxygen levels sufficiently high for the entire depth of the pond water. However, mechanical aeration is usually applied on the water surface (*e.g.*, paddlewheel aerators). Apart from being costly and energy-consuming, these aerators come with a risk of causing erosion in the pond bottom soil, when the water current is too strong. Erosion degrades embankments, makes the harvest more difficult, and damages benthic plants and animal communities, including the shrimp (Boyd, 1998). In a real pond system, also the growth of microalgae could play a critical role, as they (1) enable in situ formation of oxygen, and (2) by consuming CO_2_, they could provoke an increase in pH, which could reduce H_2_S toxicity, but increase ammonia toxicity. They can even actively contribute to an improved water quality (Huang et al., 2022). However, the direct involvement of microalgae in our model system would strongly add to the complexity of sulphide formation, because of their multi-level impact on the shrimp pond nutrient dynamics, so we eliminated the possibility for photosynthetic growth from our model by not supplying natural or artificial light. Nitrate addition could serve as an alternative electron acceptor, in competition with sulphate. However, nitrate can only temporarily control sulphide production, and when nitrate is depleted, sulphide production recovers (Schwermer et al., 2010; Torun et al., 2022; Torun et al., 2020). These limitations substantiate the importance of a lasting strategy to mitigate sulphide production in shrimp pond aquaculture systems.

In this study, 5 and 25 mg/L sodium molybdate clearly lower sulphide production in the sediment, and pushed the higher H_2_S concentration zone towards deeper sediment layers. Since the transfer of H_2_S from sediment to bulk liquid was delayed by this action, molybdate treated samples had lower concentrations of H_2_S in the bulk liquid on day 35 when peak concentrations were observed. The H_2_S concentration in the bulk liquid did fluctuate in function of time for all treatments, which can be linked to the fact that the set-up used was an open system being continuously exposed to the air inflow and disturbances created during the movement of beakers for microelectrode measurements. The re-introduction of oxygen into the bulk liquid, as confirmed by dissolved oxygen measurements, most likely re-oxidised a portion of the H_2_S. In addition, these external disturbances and the nature of the open system might have accelerated H_2_S to diffuse to the air from the bulk liquid. Alternatively, H_2_S accumulated in the bulk liquid might have reacted with ferrous iron to form iron sulphide that subsequently precipitated in the sediment.

The inhibitory concentration of molybdate and a correlation between sulphate and molybdate concentrations were shown in several studies (Biswas et al., 2009; Chen et al., 1998; Jesus et al., 2015). In our previous study, we estimated that the inhibitory concentration for 1500 mg/L sulphate present in our experimental shrimp pond model should be approximately 15 mg/L of sodium molybdate for short-term prevention of H_2_S production (Torun et al., 2022). In the current study, the molybdate was only partially reduced, both in the M5 and M25 treatments, and the production of H_2_S in the sediment and its transfer to bulk liquid could not be fully prevented. This might be due to poor diffusion of the molybdate in the deeper layer of the sediment, since in the upper layers of the sediment, there was markedly lower H_2_S concentration compared to the control treatment. The reason for molybdate having potentially lower diffusion rates than the sulphate, might be related to adsorption of molybdate on the sediment, as observed for pure quartz sand (Kögler et al., 2021). One can assume that adsorbed molybdate could not inhibit microbial sulphate reduction.

In this study, residual sulphate concentrations did not show any apparent difference between the control and molybdate treated samples, but these sulphate concentrations were measured in the bulk liquid. The H_2_S and total sulphide productions did show clear differences in the sediment itself, with higher concentrations of sulphide in the control treatment, indicating the effectiveness of molybdate to inhibit sulphate reduction. Residual sulphate in the bulk liquid remained present in all treatments, so despite the high availability of organic matter, sulphate reduction did not continue, as also observed in other studies on shrimp pond sediments (Torun et al., 2022) and other anaerobic ecosystems, such as anaerobic digestion (Lippens and De Vrieze, 2019). This apparent discrepancy was probably due to oxygen intrusion into the water column, halting sulphate reduction in the bulk liquid. Hence, sulphate reduction might have been locally interrupted in the bulk liquid, while it continued in the deeper layers of the sediment.

Overall, it is clear that the biogeochemical sulphur cycle in such a pond system involves various processes, i.e., sulphate reduction, sulphide/sulphur (re-)oxidation, precipitation of metal sulphides, and production of polysulphides. Due to the nature of the open air system (as is the case in real pond systems) in the current study, with the possibility of H_2_S escaping, it is not possible to make accurate sulphur mass balances.

### 4.2. Molybdate treatment changes the absolute abundance of sulphate reducing bacteria

When molybdate is provided in the presence of SRB, ATP sulfurylase uses molybdate (instead of sulphate) and ATP to produce an unstable molecule equivalent to adenosine 5′-phosphosulfate (APS) that cannot be used as electron acceptor (Biswas et al., 2009). Under molybdate excess, some studies indicated that SRB growth could be supressed altogether. Kögler et al. (2021) showed that there were no SRB specific *dsR* genes isolated when molybdate was continuously injected into sandpacks with residual oil in an oil reservoir. Nair et al. (2015) reported that molybdate concentrations ranging between 50 and 150 µM increased the doubling time of *Desulfovibrio alaskensis* G20, and 500 µM molybdate completely inhibited its cellular growth. In the current study, molybdate was provided only once at lower concentrations than the concentrations mentioned in the literature, but even such lower concentrations of molybdate showed a promising impact towards decreasing sulphide concentrations in the bulk liquid.

A higher absolute abundance of the phylum Delsufobacterota, containing several SRB, was detected in molybdate treated samples, despite the fact that mitigation of sulphide production was observed. A similar trend was observed in an earlier study (Tenti et al., 2019), where SRB counts in all samples from a lab-scale anaerobic digester, were similar with or without molybdate, when molybdate concentration was lower than 1.2 mM. In this study, the detection of SRB through 16S rRNA gene amplicon sequencing showed a relative increase of SRB, but did not provide any information on their activity or absolute abundance. Hence, these relative abundances were combined with the absolute cell counts, as obtained through flow cytometry analysis to estimate absolute cell counts of the Desulfobacterota phylum. Such an approach has been successfully applied in other ecosystems, and can be considered an established, reliable way of quantifying microorganisms in environmental samples (Barr et al., 2021; Ou et al., 2017; Props et al., 2017). An overall increasing trend in time was observed for the Desulfobacterota phylum, including the samples treated with molybdate. The reason for the higher absolute abundance of SRB, despite lower H_2_S production, might be related to (partial) inactivation of enzymes involved in sulphate reduction. In the study of Nair et al. (2015) on growth and morphology of *Desulfovibrio alaskensis* G20, at least three important enzymes that play a crucial role in energy production (alcohol dehydrogenase, pyruvate carboxylase, tungsten formylmethanofuran dehydrogenase) showed downregulation or repression in the presence of elevated molybdate concentrations. In the current study, molybdate treatment at 25 mg/L showed an overall significantly different bacterial community composition compared to the control without molybdate. The increase in SRB was unexpected, yet, next to sulphate reduction, SRB can also carry out hydrogenic and/or acetogenic metabolisms. Hence, in the absence of sulphate, many SRB can ferment organic acids or alcohol, producing hydrogen acetate or carbon dioxide (Plugge et al., 2011; Zhang et al., 2022). The growth of *Desulfovibrio* on lactate was reported in the absence of sulphate, in syntrophy with a methanogen (Bryant et al., 1977), and the growth of the Delsufovibrionaceae family was also detected in this study. Overall, a combination of reduced H_2_S toxicity and the shift in the energy production metabolism appeared to have increased the relative abundance of SRB in this study.

## 5. Conclusions

We showed that molybdate could be an effective mitigation agent against sulphide accumulation in shrimp ponds, since it can be applied in a single dose, and at relatively low concentrations. Although, sulphide production could not be avoided completely, and only a temporal effect could be obtained, molybdate reduced H_2_S production in the sediment, and delayed its transfer to the water column by pushing the sulphide production zone towards deeper sediment layers. Molybdate induced a higher absolute abundance of Desulfobacterota, but this was not reflected in increased sulphide formation. Overall, molybdate **has the potential to** serve as a more environmentally friendly option compared to other conventional strategies to mitigate sulphide production in shrimp pond systems.

## Acknowledgments

Special thanks go to the Laboratory of Aquaculture and Artemia Reference Center, Ghent University, for providing shrimp feed (Crevetec Grower 2) and faeces. We would like to thank Koenraad Maréchal and Dirk Raes from Agentschap Natuur & Bos (Belgium) for their help during collection of sediment samples, Tim Lacoere for his assistance with the molecular analyses and Sam Decroo for his assistance with the flow cytometry. The authors also kindly acknowledge Peter Goethals, Peter Bossier, Marleen De Troch and Laurine Burdorf for carefully reading the manuscript.

## Data, script, code and supplementary material

The datasets generated and R scripts used during this research are included in this article, its supplementary information, and were submitted to the Zenodo repository (https://zenodo.org/doi/10.5281/zenodo.10149234). The raw fastq files that served as a basis for the bacterial community analysis were deposited in the National Center for Biotechnology Information (NCBI) database (Accession number SRP326102).

