## Supplementary file 1 for "Molybdate delays sulphide formation in the sediment and transfer to the bulk liquid in a model shrimp pond"

Journal: bioRxiv

Title: Long-term sulphide mitigation through molybdate at shrimp pond bottoms

### S1. Shrimp feeding tables

**Table S1** Daily feeding table for the shrimp post-larvae (PL) stocked at 50 shrimp per $m^3$ density

| Day of culture (DOC) | PL stage | PL size (mg) | Daily feeding rate (% of PL mass) | Required feed ($g\ m^{-3}\ d^{-1}$) |
|---|---|---|---|---|
| 0 | 10 | 6 | 34 | 0.102 |
| 1 | 11 | 7 | 33 | 0.107 |
| 2 | 12 | 7 | 32 | 0.112 |
| 3 | 13 | 11 | 31 | 0.171 |
| 4 | 14 | 15 | 30 | 0.225 |
| 5 | 15 | 20 | 29 | 0.29 |
| 6 | 16 | 25 | 28 | 0.35 |
| 7 | 17 | 29 | 28 | 0.406 |
| 8 | 18 | 33 | 27 | 0.446 |
| 9 | 19 | 38 | 27 | 0.513 |
| 10 | 20 | 45 | 27 | 0.608 |
| 11 | 21 | 60 | 26 | 0.780 |
| 12 | 22 | 80 | 26 | 1.040 |
| 13 | 23 | 100 | 26 | 1.300 |
| 14 | 24 | 105 | 25 | 1.312 |
| 15 | 25 | 110 | 25 | 1.375 |
| 16 | 26 | 135 | 22 | 1.485 |
| 17 | 27 | 155 | 20 | 1.550 |
| 18 | 28 | 185 | 20 | 1.850 |
| 19 | 29 | 215 | 19 | 2.043 |
| 20 | 30 | 245 | 19 | 2.328 |
| 21 | 31 | 275 | 18 | 2.475 |
| 22 | 32 | 310 | 18 | 2.790 |
| 23 | 33 | 345 | 18 | 3.105 |
| 24 | 34 | 400 | 17 | 3.400 |
| 25 | 35 | 465 | 17 | 3.953 |
| 26 | 36 | 550 | 17 | 4.675 |
| 27 | 37 | 640 | 16 | 5.120 |
| 28 | 38 | 695 | 16 | 5.560 |
| 29 | 39 | 750 | 16 | 6.000 |
| 30 | 40 | 835 | 15 | 6.263 |
| 31 | 41 | 920 | 15 | 6.900 |
| 32 | 42 | 1025 | 15 | 7.688 |
| 33 | 43 | 1130 | 14 | 7.910 |
| 34 | 44 | 1240 | 14 | 8.680 |
| 35 | 45 | 1360 | 14 | 9.520 |
| 36 | 46 | 1480 | 14 | 1.360 |
| 37 | 47 | 1600 | 13 | 10.400 |

| | | | | |
|---|---|---|---|---|
| 38 | 48 | 1720 | 13 | 11.180 |
| 39 | 49 | 1850 | 13 | 12.025 |
| 40 | 50 | 1970 | 13 | 12.805 |
| 41 | 51 | 2100 | 12 | 12.600 |
| 42 | 52 | 2250 | 12 | 13.500 |
| 43 | 53 | 2340 | 12 | 14.040 |
| 44 | 54 | 2434 | 12 | 14.602 |
| 45 | 55 | 2531 | 11 | 13.920 |
| 46 | 56 | 2632 | 11 | 14.477 |
| 47 | 57 | 2737 | 11 | 15.056 |
| 48 | 58 | 2847 | 11 | 15.658 |
| 49 | 59 | 2961 | 10 | 14.804 |
| 50 | 60 | 3079 | 10 | 15.396 |
| 51 | 61 | 3202 | 10 | 16.012 |
| 52 | 62 | 3331 | 10 | 16.658 |
| 53 | 63 | 3464 | 10 | 17.319 |
| 54 | 64 | 3602 | 9 | 16.211 |
| 55 | 65 | 3746 | 9 | 16.859 |
| 56 | 66 | 3896 | 9 | 17.534 |
| 57 | 67 | 4052 | 9 | 18.235 |
| 58 | 68 | 4214 | 9 | 18.964 |
| 59 | 69 | 4383 | 8 | 17.531 |
| 60 | 70 | 4558 | 8 | 18.232 |
| 61 | 71 | 4740 | 8 | 18.961 |
| 62 | 72 | 4930 | 8 | 19.720 |
| 63 | 73 | 5127 | 8 | 20.509 |
| 64 | 74 | 5332 | 7 | 18.663 |
| 65 | 75 | 5546 | 7 | 19.410 |
| 66 | 76 | 5767 | 7 | 20.186 |
| 67 | 77 | 5998 | 7 | 20.993 |
| 68 | 78 | 6238 | 7 | 21.833 |
| 69 | 79 | 6488 | 6 | 19.463 |
| 70 | 80 | 6747 | 6 | 20.241 |
| 71 | 81 | 7017 | 6 | 21.051 |
| 72 | 82 | 7298 | 6 | 21.893 |
| 73 | 83 | 7590 | 6 | 22.769 |
| 74 | 84 | 7893 | 5 | 19.733 |
| 75 | 85 | 8209 | 5 | 20.522 |
| 76 | 86 | 8537 | 5 | 21.343 |
| 77 | 87 | 8879 | 5 | 22.197 |
| 78 | 88 | 9234 | 5 | 23.085 |
| 79 | 89 | 9603 | 4 | 19.206 |
| 80 | 90 | 9987 | 4 | 19.975 |
| 81 | 91 | 10387 | 4 | 20.774 |
| 82 | 92 | 10802 | 4 | 21.605 |

| | | | | |
|---|---|---|---|---|
| 83 | 93 | 11234 | 4 | 22.469 |
| 84 | 94 | 11684 | 4.5 | 26.288 |
| 85 | 95 | 12151 | 4.5 | 27.340 |
| 86 | 96 | 12637 | 4.5 | 28.433 |
| 87 | 97 | 13143 | 4.5 | 29.570 |
| 88 | 98 | 13668 | 4.5 | 30.754 |
| 89 | 99 | 14215 | 3.5 | 24.876 |
| 90 | 100 | 14784 | 3.5 | 25.871 |

#### S2. Organic waste supplementation table

**Table S2** The amount of organic waste supplemented at the different periods of the experiment. The experiment was started with an initial waste load of DOC (day of culture) 30, and additional feed and faeces were supplemented every 2-3 days to simulate the waste accumulation. The amount of feed and faeces were increased every 15 days to account for the increasing uneaten feed and faeces accumulation on the pond bottom with the increasing size of shrimp, due to growth. The volume of faeces supplemented was determined based on its dry matter content (4.32%).

| Period (Waste load/ experiment) | Feed supplement (mg/day) | Faeces supplement (mL/day) |
|---|---|---|
| DOC 30/ Day 0 (initial) | 27 | 0.82 |
| DOC 30-45/ Day 0-15 | 4.84 | 0.15 |
| DOC 45-60/ Day 15-30 | 7.26 | 0.22 |
| DOC 60-75/ Day 30-45 | 8.92 | 0.27 |
| DOC 75-90/ Day 45-60 | 10.61 | 0.32 |

#### S3. Amplicon sequencing and data processing

S3.1. Amplicon sequencing

The primers 341F (5’- CCTACGGGNGGCWGCAG) and 785Rmod (5’- GACTACHVGGGTATCTAAKCC) that target the V3-V4 region of the 16S rRNA gene (Klindworth et al., 2013), with an extra wobble position in the reverse primer to make it more universal, were used to target total bacteria. The DNA extracts had a minimum DNA concentration of 10 ng $\mu L^{-1}$, and were free of RNA, which was validated with agarose gel electrophoresis. A 2-step PCR protocol was used, and 10-25 ng of DNA was used as template for the first PCR run with a total volume of 50 µL, using the 341F and the 785Rmod primers with Illumina adaptor sequences. The first PCR run contained 25 cycles, with an annealing temperature of 55 °C. The PCR products were purified using Ampure XP beads according to the manufacturer’s instructions, and their size checked on a Fragment analyser (Advanced Analytical Technologies, Inc., Ames, Iowa, USA), and quantified by fluorometric analysis. The purified PCR products were subjected to a 2$^{nd}$ PCR run that contained 6 cycles with an annealing temperature of 55 °C, using sample-specific barcoded primers (Nextera XT index kit, Illumina). The PCR products were again purified using Ampure XP beads according to the manufacturer’s instructions, and their size checked on a Fragment analyser, and quantified. Next, multiplexing, clustering and sequencing was carried out on an Illumina MiSeq with the paired-end (2×) 300 bp protocol and indexing. The sequencing run was analysed *via* the Illumina CASAVA pipeline (v1.8.3) in which demultiplexing was based on sample-specific barcodes. The raw sequencing data were processed, removing sequence reads of too low quality (only "passing filter" reads were selected), and discarding reads containing adaptor sequences or PhiX control with an in-house filtering protocol. A quality assessment on the remaining reads was performed using the FASTQC quality control tool version 0.10.0.

S3.2. Data processing

The Mothur software package (v.1.44.3), and guidelines developed by Schloss et al. (2009) were used to process the raw Illumina data on a x86_64-pc-linux-gnu (64-bit) system. The forward and reverse reads were assembled into contigs by a heuristic approach, taking the Phred quality scores into account. Ambiguous contigs or with unsatisfying overlap were removed, and the remaining sequences were aligned to the mothur formatted silva seed v138 database. Those sequences not aligning within the region targeted by the primer set or sequences with homopolymer stretches with a length > 12 bp were removed. The sequences were pre-clustered, allowing mismatch for every 100 bp of sequence. Chimeric sequences were removed with VSEARCH (Rognes et al., 2016). Classification of the sequences was carried out by a naïve Bayesian classifier (Wang et al., 2007), using the RDP 16S rRNA gene training set, release 16, with an 85% cut-off for the pseudobootstrap confidence score. Taxa that were annotated as Chloroplast, Mitochondria, unknown, Archaea or Eukarya at the kingdom level were excluded. Sequences were clustered into OTUs (operational taxonomic units) with an average linkage, and at a 97% sequence identity, using the OptiClust method (Westcott and Schloss, 2017). If sequences could not be classified at the (super)Kingdom level, they were removed. Representative sequences were picked for each OTU as the most abundant sequence within that OTU.

#### S4. Rarefaction curves

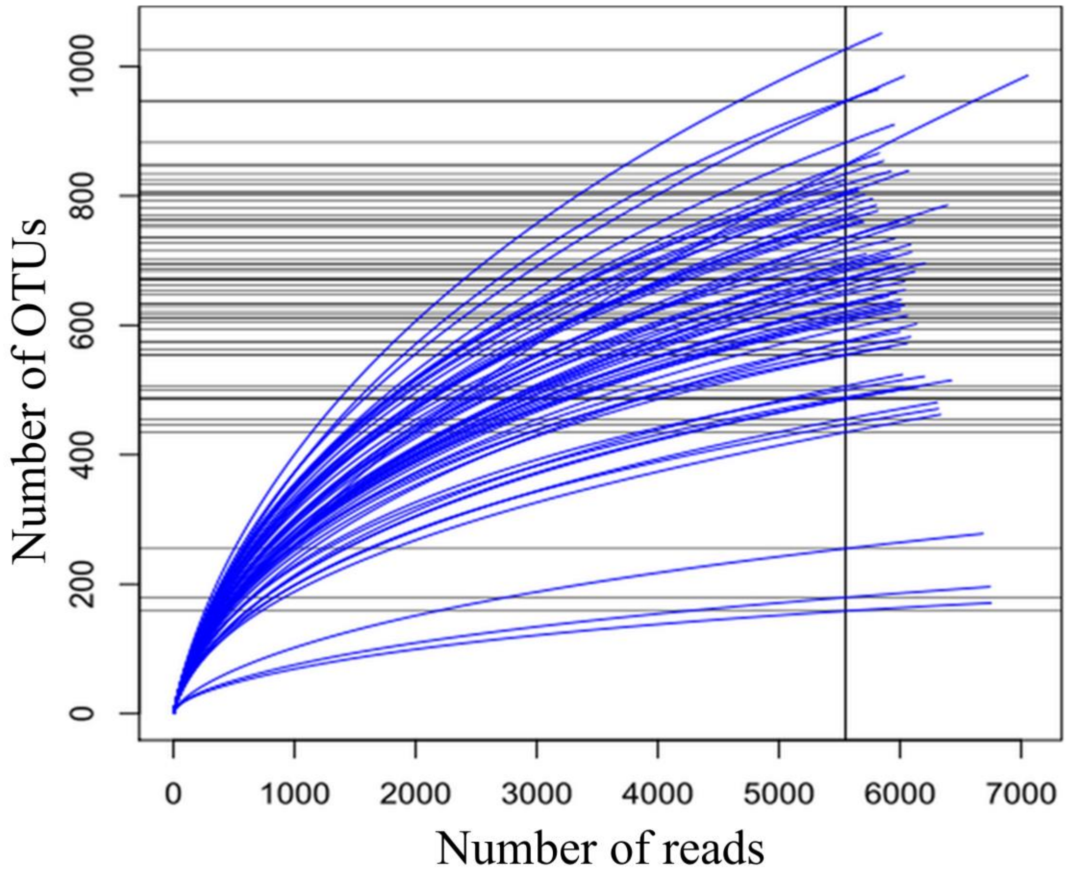

**Figure S**1 Rarefaction curves indicating the number of resolved OTUs against sampling depth of each of the samples.

#### S5. Additional bulk measurements

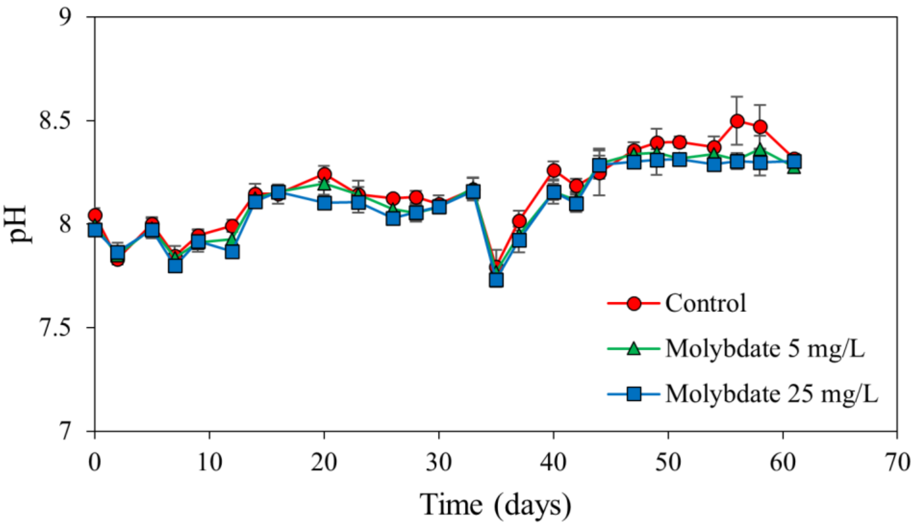

**Figure S2** The bulk liquid pH in the control treatment, molybdate treatment at 5 mg/L (M5) and molybdate treatment at 25 mg/L (M25). Values represent averages of biological triplicates, and error bars represent the standard deviation.

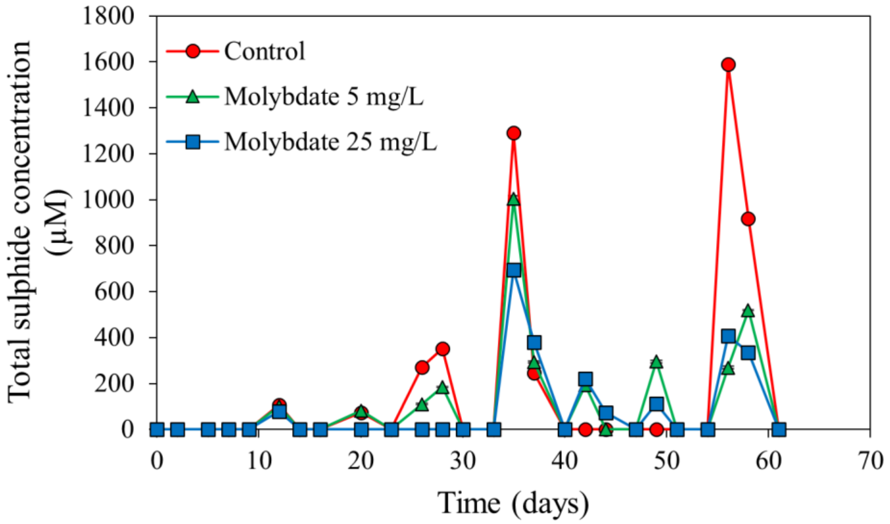

**Figure S3** The bulk liquid total sulphide concentration in the control treatment, molybdate treatment at 5 mg/L (M5) and molybdate treatment at 25 mg/L (M25). Values represent averages of biological triplicates, and error bars represent the standard deviation.

#### S6. Additional sediment profiles

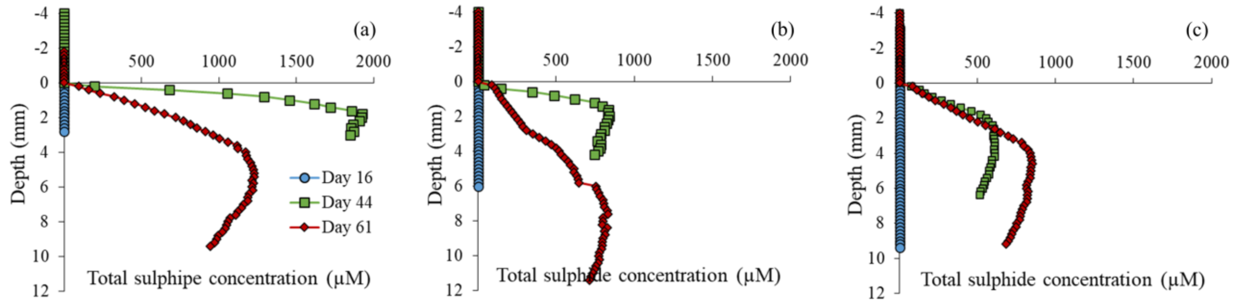

**Figure S4** The total sulphide depth profiles for the (a) control treatment, (b) molybdate treatment at 5 mg/L (M5) and (c) molybdate treatment at 25 mg/L (M25). Values represent averages of biological triplicates, error bars are omitted to maintain the visibility of the graphs. Zero depth equals to the sediment-water interface. Because of technical problems with the microelectrode, data from day 30 are not included.

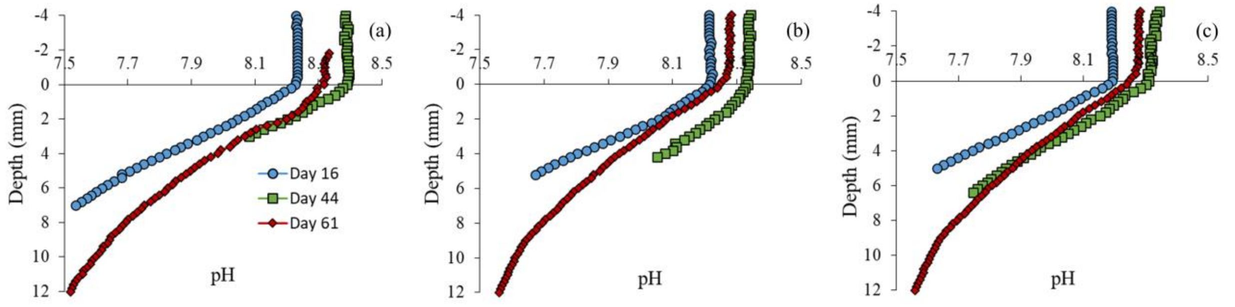

**Figure S5** The pH depth profiles for the (a) control treatment, (b) molybdate treatment at 5 mg/L (M5) and (c) molybdate treatment at 25 mg/L (M25). Values represent averages of biological triplicates, error bars are omitted to maintain the visibility of the graphs. Zero depth equals to the sediment-water interface. Because of technical problems with the microelectrode, data from day 30 are not included.

### S7. Heatmap at the family level

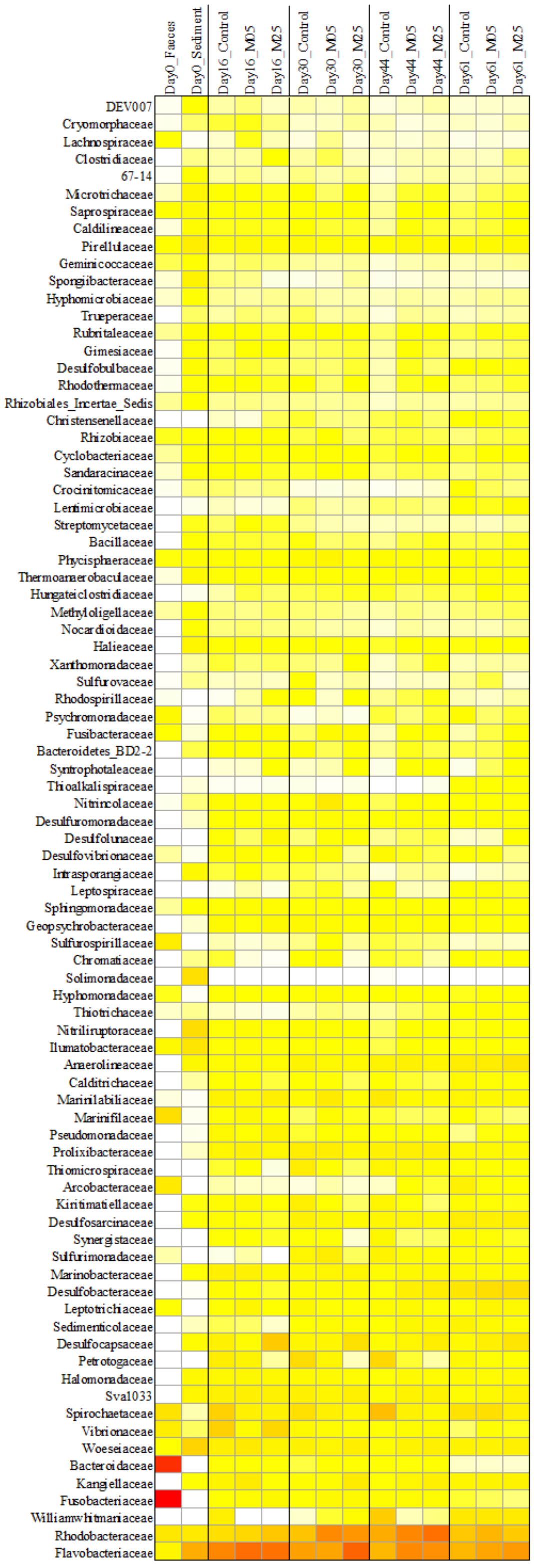

**Figure S6** Heatmap showing the relative abundance of the bacterial community at the family level in the faeces, the sediment and the different treatments on day 16, 30, 44 and 61. Weighted average values of the biological replicates are presented. The colour scale ranges from 0 (white) to 35% (red) relative abundance.
